# A winner in the Anthropocene: changing host plant distribution explains geographic range expansion in the gulf fritillary butterfly

**DOI:** 10.1101/754010

**Authors:** Christopher A. Halsch, Arthur M. Shapiro, James H. Thorne, David P. Waetjen, Matthew L. Forister

## Abstract

1. The changing climate is altering the geographic distributions of species around the world with consequences for population dynamics, resulting in winners and losers in the Anthropocene.
2. *Agraulis vanillae*, the gulf fritillary butterfly, has expanded its range in the past one hundred years in the western United States. We combine time series analysis with species distribution modeling to investigate factors limiting the distribution of *A. vanillae* and to predict future shifts under warming scenarios.
3. In the western US, where we have time series and geographic data, urban development has a positive influence (the host plant is an ornamental in gardens), being associated with year of colonization. Colonization was also associated to a lesser extent with winter maximum temperatures, while a negative impact of minimum temperatures and precipitation was apparent on population growth rates.
4. Across the country, the butterfly is primarily limited by host availability and positively affected by human presence. Perhaps counter-intuitively for a largely tropical ectotherm, current effects of a warming climate (in the years post-colonization) are either negative (on population growth rate) or indirect, likely mediated through availability of areas that can support the host.
5. Under future climate scenarios, conditions are predicted to become more suitable for *Passiflora* in many urban areas, which would likely result in further expansion of *A. vanillae* during the dispersive season. These results illustrate the value of combining time series with spatial modeling to understand and predict shifting geographic ranges in the Anthropocene.

## Introduction

Recent climate change has had numerous consequences for species around the world, including shifts in geographic distribution (Chen et al., 2011). In some cases, ranges are expanding, while for many others geographic ranges are shifting or contracting (Parmesan, 2006). Ectotherms, including butterflies and other insects, are particularly sensitive to changes in the climate and have often been exemplar species for the study of these issues (Parmesan et al., 1999; Warren et al., 2001). Recently, attention has been paid not only to changes in geographic ranges, but also to declines in insect populations around the world, which are driven by a combination of habitat loss, pesticide use, climate change, and other factors (Hallmann et al., 2017; Lister & Garcia, 2018; Salcido et al., 2019; Sanchez-Bayo & Wyckhuys, 2019; Wepprich et al., 2019). We can expect these factors will have different impacts on different species, and even that some species will be “winners” under altered conditions (McKinney & Lockwood, 1999). Identifying successful species and the reason for their success in the face of change is important for understanding the potential of individual species and ecosystems to persist and thrive in future climates. In particular, understanding how aspects of global change negatively impact some species, while benefiting others, will improve our ability to predict future species assemblages. One example of a butterfly that appears to be benefitting from anthropogenic influence is the gulf fritillary (*Agraulis vanillae)*, which has recently expanded its range in the western United States (Shapiro, 2007). In this study we seek to better understand the drivers underlying this expansion using a combination of spatial data and long-term population records.

*Agraulis vanillae* is a neotropical butterfly associated with riparian and weedy/disturbed habitats (Shapiro, 2009). Over its entire distribution, from temperate North America to temperate South America, there are eight identified sub-species. Previous work has demonstrated genetic divergence between North American and South American lineages (Runquist et al., 2012). In the United States, *A. vanillae* is multi-voltine and in some areas flies almost all year, however diapause has been observed in both the larval and pupal stages in Florida (Sourakov, 2008). The butterfly is sensitive to frost, which can be lethal to all life stages (Shapiro, 2007). Eastern populations are known to undergo northward dispersal in the spring (Walker, 1991), with sightings as far north as North Dakota and New York (Scott, 1986). These life history traits raise the possibility that the range of *A. vanillae* may be limited by low overwintering temperatures, thus milder winters could reduce the risk of extinction along the northern range margin and explain the success of this butterfly.

*Agraulis vanillae* utilizes most plants from the genus *Passiflora* as hosts (May, 1992). The two most common species in the United States are *Passiflora incarnata* and *Passiflora lutea*, both of which are found across much of the southeastern United States (Gremillion, 1989). *Passiflora* prefers well-drained soils and is often found in disturbed sites. In the west, *Passiflora* is not present in natural areas, however various species have been introduced to urban areas as ornamentals (Graves & Shapiro, 2003). Winter low temperatures likely limit the distributions of wild populations, however survival can be improved by active management in cultivated populations (McGuire, 1999). In California, the introduction of *Passiflora* and its association with *A. vanillae*, are well documented. In Southern California, these species have been associated since 1875. It was first sighted in San Francisco as early as 1908, however it did not permanently establish until 1955 (Powell, 2000). In the 1960’s and 1970’s the butterfly was seen in Sacramento, but was extirpated and has only recently reestablished in the region. The presence of *Passiflora* offers another, non-mutually exclusive, explanation for the success of *A. vanillae*. It is possible that *A. vanillae* is currently not limited by temperature, but instead by the distribution of *Passiflora.* As this plant expands due to cultivation, so does the gulf fritillary.

In this study, we address the following questions. First, using data from a long-term observational study, we ask if climate or urban development better explain the establishment and success of the butterfly in recent years in the Sacramento Valley. Second, using citizen science observational data, we ask if the current distribution of the butterfly in the continental United States is better explained by host plant or climate limitation. Finally, using species distribution modeling, we ask if the butterfly is likely to continue to expand its distribution under different climate change scenarios.

## Materials and methods

### Sacramento Valley time series data

Observational data were collected every other week by a single observer (AMS) across five sites in the Sacramento Valley. Count data of individual butterflies at these five sites have been collected since 1999 and presence/absence data have been collected since the 1970’s or 1980’s, depending on the site. At these five low elevation sites, data are recorded year round. Site descriptions and additional details have been reported elsewhere (Forister et al., 2010). *Agraulis vanillae* did not consistently appear at any of these five sites until 2001 and did not appear at every site until 2012. Climate data in California were derived from 270m grid climate maps of monthly and annual values for minimum and maximum temperature and precipitation (Flint & Flint 2012; Flint et al. 2013; Thorne et al. 2015). We extracted the values for grid cells that overlapped with each of the sample sites in the Sacramento Valley and averaged the values for each monthly variable for each year. We calculated seasonal variables by further averaging monthly values to season and converting to water year (the start of September through the end of August).

### Sacramento Valley statistical analysis

We approached the analysis of times series data in two phases. First, we used annual presence/absence data to examine colonization, attempting to model the difference between years in which the butterfly was absent across our focal sites and years in which it was present (spanning 1984 through 2018). Specifically, random forest regression was used with presence at a site in a given year as the response variable and percent urban land cover (at a county level), seasonal means of minimum temperature, seasonal means of maximum temperature, and seasonal means of precipitation as covariates. A total of 500,000 trees were made with a node size of 5. Variable importance was determined by examining the increased mean squared error of the model when each variable was randomly permuted. The most influential variables identified by random forest analysis were moved forward into a Bayesian hierarchical model. While the random forest is useful for judging the potential importance of a large number of variables, including some that are highly correlated, the Bayesian model allows us to estimate coefficients and associated uncertainty in a hierarchical framework (simultaneously within and across sites). Following a previous model used for data from these study sites (Nice et al, 2019), presence was modeled both at the individual site level and at a higher level across all sites using a Bernoulli distribution. Uninformative priors were used for means and variance, with means drawn from normal distribution (mu = 0, tau = 0.01) and variances drawn from a gamma distribution (r = 0.01, lambda = 0.01). The Bayesian model was comprised of four chains each run for 5,000,000 iterations with an adaptive phase of 500,000 iterations.

As a second phase, we examined annual population dynamics post-colonization at the same focal sites, using individual survey count data summarized by year and transformed into population growth rates. Population growth was calculated as the natural log of the current year’s total count divided by the previous year’s total count (Sibly & Hone, 2002). To determine the most influential climate variables, population growth in a given year was then modeled using a random forest regression. Covariates in the model included abundance in the previous year, seasonal means of minimum monthly temperature, seasonal means of maximum monthly temperature, seasonal means of precipitation, and these same variables lagged by one year to allow in particular for effects mediated through host plants. Again, a total of 500,000 trees with a node size of 5 was used. Variable importance was determined by examining the increased mean squared error of the model following permutation of each variable, and this was done both within and among sites. Like the colonization analysis, the most influential variables identified by random forest analysis were moved forward into a Bayesian hierarchical model in which population growth was modeled both at the individual site level and at a higher level across all sites using a normal distribution. Means of covariates were drawn from an uninformed normal distribution (mu = 0, tau = 0.01) and variances drawn from an uninformed gamma distribution (r = 0.01, lambda = 0.01). This model was comprised of four chains each run for 100,000 iterations with an adaptive phase of 10,000 iterations. All analyses were conducted using the randomForest (RColorBrewer & Liaw, 2018) and jagsUI (Kellner, 2019) packages in R Studio.

### National data

For US-wide spatial analyses, geo-referenced data points for both *A. vanillae* and *Passiflora* were acquired from “research grade” observations on inaturalist. Additional observations of *Passiflora* were obtained from Cal flora and additional observations of *A. vanillae* from the Butterflies and Moths of North America. Only observations with an uncertainty under 1km were used for analysis. Both *Passiflora* and *A. vanillae* are distinct and identification is likely not a concern, however a random subset of 100 observations with photos were checked and all were found to be correct IDs. Current climate data and future projections were obtained from WorldClim (Hijmans et al., 2005). A human population density raster was obtained from the Socioeconomic Data and Applications Center, which used data from the 2010 census (Center for International Earth Science Information Network, 2018). All raster layers were cropped to only include the 48 contiguous states, although *A. vanillae* is also an exotic in Hawaii. Finally, *A. vanillae* points were separated based on being from the dispersal season or before. Points from January to March were labeled as pre-dispersal, which is earlier than the earliest observed spring migrant from a study of this movement in Florida (Walker, 1991). In this paper, dispersal will refer to the maximum distribution that the butterfly achieves during the year.

### National statistical analysis

Species distribution models were built for both *Passiflora* and *Agraulis vanillae.* All host plant models were built at the genus level, however *Passiflora* species known not to be host plants were excluded. The western and eastern distributions were modeled both separately and together, to allow for the possibility of different factors affecting range limits in the different regions. For all models, we used the maxent algorithm, which models presence only data by comparing observations with random background points. For every model, 10,000 random background points were taken within the continental United States. *Passiflora* was modeled using temperature, mean precipitation, and human population density as covariates. Models were built and evaluated using mean temperature in the coldest month, mean annual temperature, and max temperature in the warmest month as the temperature variables. The best performing host plant model was later used as part of the butterfly spatial model. Since the host plant, especially in the western United States, is found almost exclusively in urban environments, human population density was used as a proxy for urban cultivation of the plant. For *A. vanillae*, both the overwintering distribution and dispersal distributions were modeled. The overwintering distribution was modeled using the *Passiflora* distribution model and temperature variables. The dispersal distribution was similarly modeled using the *Passiflora* distribution model and temperature as covariates. As with *Passiflora* analyses, various temperature variables were used for model building and comparison, and only the highest performing model for both overwinter and dispersal distributions were used for inference and projection. The models were trained on 70% of the data and tested with the remaining 30%. For all models, data were thinned in order to reduce overfitting due to sampling bias, which was done by creating a grid that was overlaid onto the study area. Individual grid cells were approximately 50 km^2^ and if more than five observations were located within a single grid, only a random subset of five was kept for analysis. If a grid cell had less than five observations all were kept. A sensitivity analysis was also performed to examine the impacts of the thinning method. All data used in models were thinned substantially, often by over fifty percent (Table S1). Model evaluation was performed by examining the AUC scores and omission error rates of both the real model and 1000 permuted null models. Methods and code for null model permutation are described by Bohl et al. (2019), but briefly, observations from the real model are randomly moved around the study area and compared to the real model using the same covariates and testing data. All analyses were performed in R Studio using the dismo package (Hijmans et al., 2013).

## Results

### Time Series

For the first twenty-five years of the time series, *Agraulis vanillae* only appeared as an occasional visitor, however beginning in 2001 it became a frequent visitor to all sites across the Sacramento Valley. This rise in the presence of *A. vanillae* occurred during a time of rising temperature and increasing urban development in the area (fig. 1). The random forest model attributed high importance to winter maximum temperatures and percent urban land cover in predicting presence at a site (fig. 2). Both maximum temperature and urban land cover have increasing trends over time, especially land cover, which is highly correlated with year (correlation coefficients for year and land cover range from 0.973 in Solano county to 0.989 in Yolo county). For the Bayesian model, the model successfully converged (as judged by visual inspection of posterior probability distributions) at both the individual site level and at the higher across site level. The Bayesian model confirms that both maximum winter temperatures and development are positively associated with colonization at the higher across site level. Specifically, the probability that maximum temperature has a greater than zero effect is 0.98 and the probability that urban development has a greater than zero effect is 0.88. The coefficient estimate for urban development shows high uncertainty for any particular value, however the posterior distribution is almost entirely greater than the maximum temperature posterior, thus there is support for a stronger effect of urbanization.

**Figure 1.**
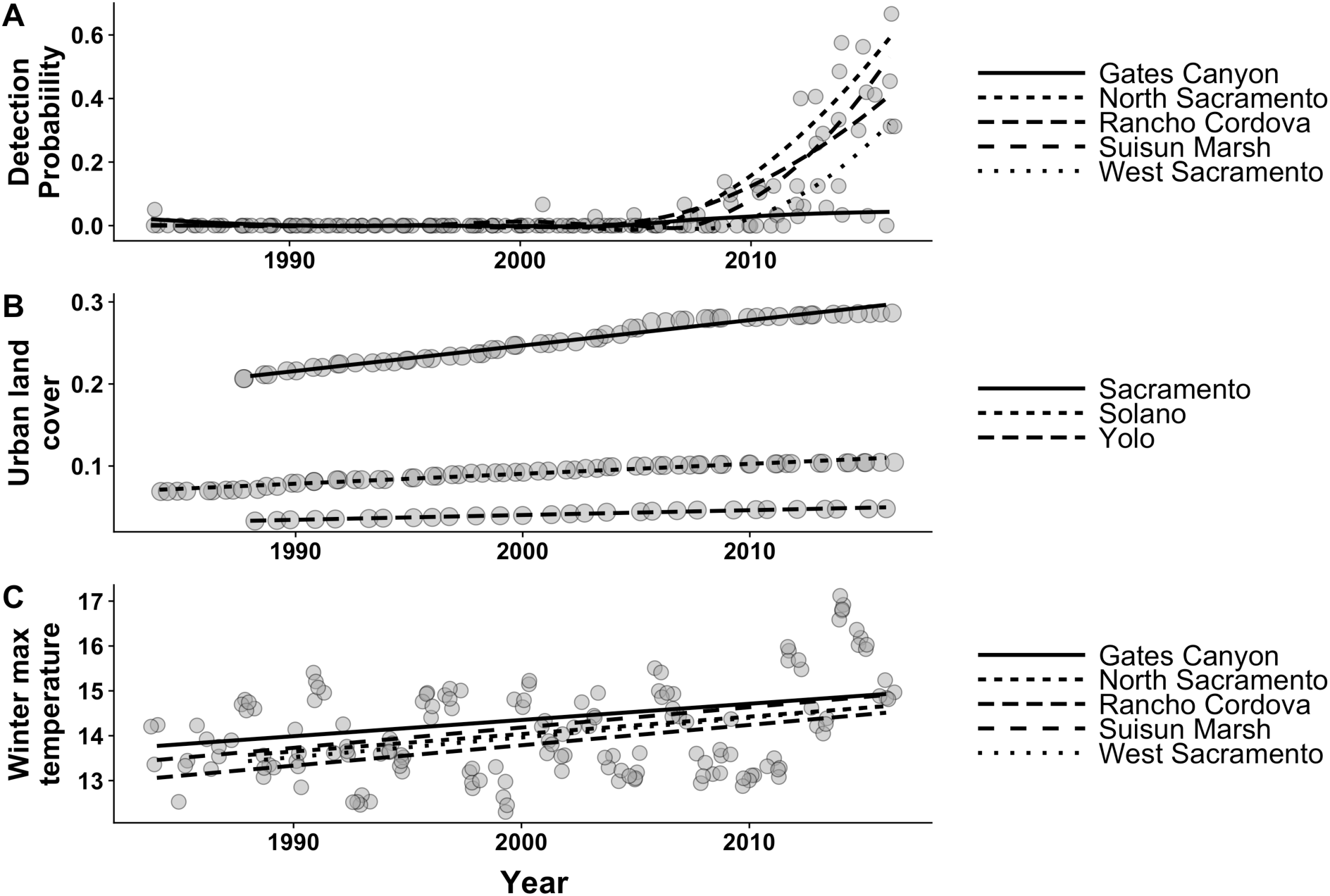
(a) Change in detection probability (the ratio of days observed to total visits) over time across all sites. (b) Annual ratio of urban land cover to total land cover at a county level for the three counties containing long-term study sites: North Sacramento and Rancho Cordova are in Sacramento County; Suisun Marsh and Gates Canyon are in Solano County. (c) Mean monthly maximum winter temperature over time.

**Figure 2.**
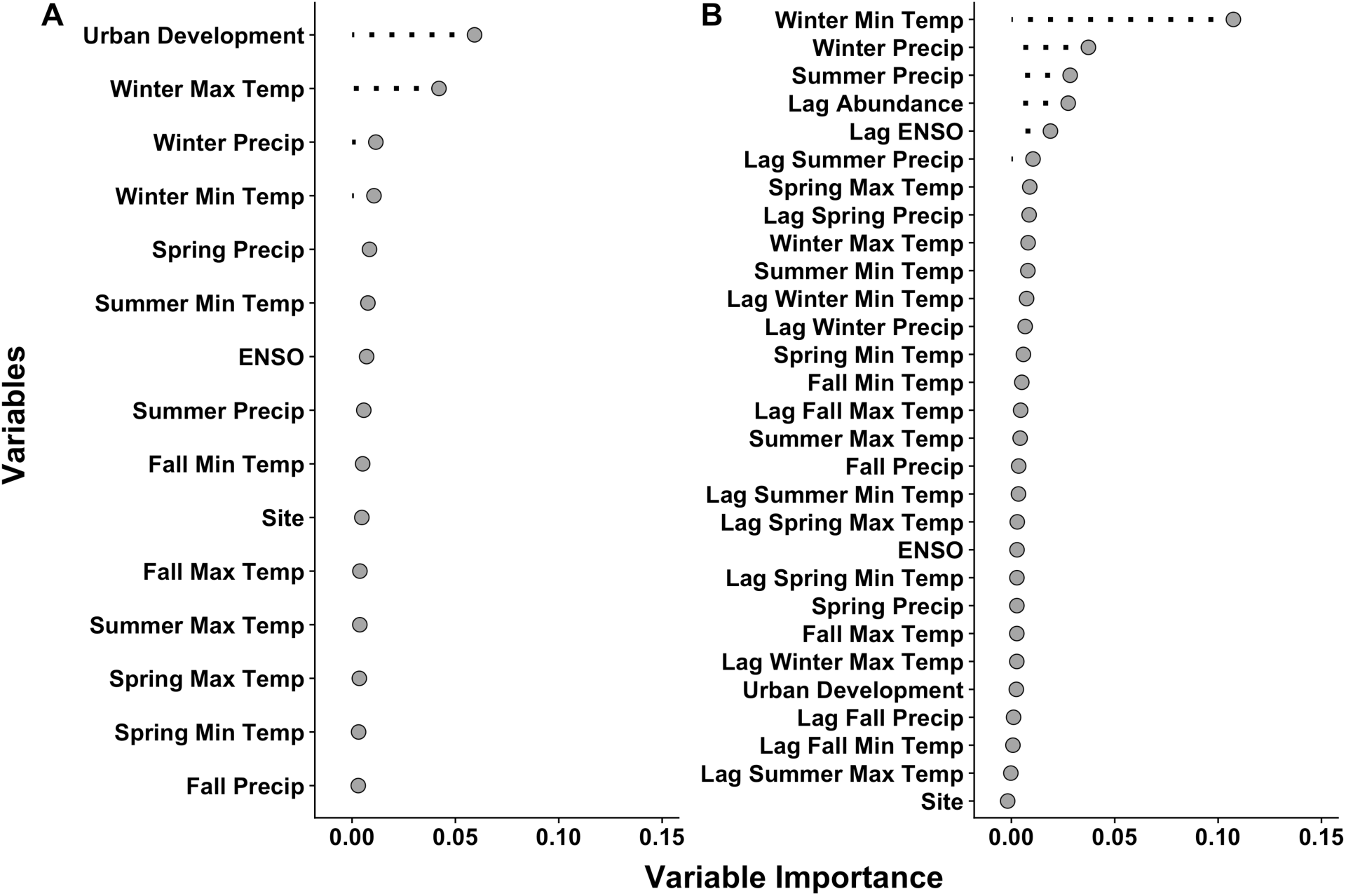
(a) Variable importance of model covariates in predicting the presence of *A. vanillae* at a site in the Sacramento Valley over time. (b) Variable importance of model covariates in predicting the annual population growth after establishment.

For annual population dynamics (represented by the natural log of the current to previous population density), the random forest analysis attributed high importance to abundance in the previous year, winter minimum temperature in the current year, winter precipitation in the current year, and summer precipitation in the current year for predicting population growth (fig. 2, fig. S1). Urbanization, while one of the covariates in the model, was not found to be important for population growth rates. Coefficients in the Bayesian model for population growth converged at both the across site and individual site level. Previous year’s abundance, winter minimum temperature, and winter precipitation all had negative effects on population growth. The model is confident in the negative impacts of previous year’s abundance, winter minimum temperature, and winter precipitation (fig. 3). Specifically, the probability that previous year’s abundance has a negative effect is 0.84, the probability that winter minimum temperature has a negative effect is 0.80, and the probability that winter precipitation has an effect is 0.88. There does not appear to be a strong effect of summer precipitation in the Bayesian hierarchical regression, despite the importance attributed to it in the random forest. All three variables have approximately equal estimated effect sizes. At the individual site level, there is variation in estimated effects, however negative density dependence is observed at all sites. Winter climate is also important at all sites, however some sites have higher estimated impacts of winter precipitation while others more heavily weight winter minimum temperatures (fig. S4).

**Figure 3.**
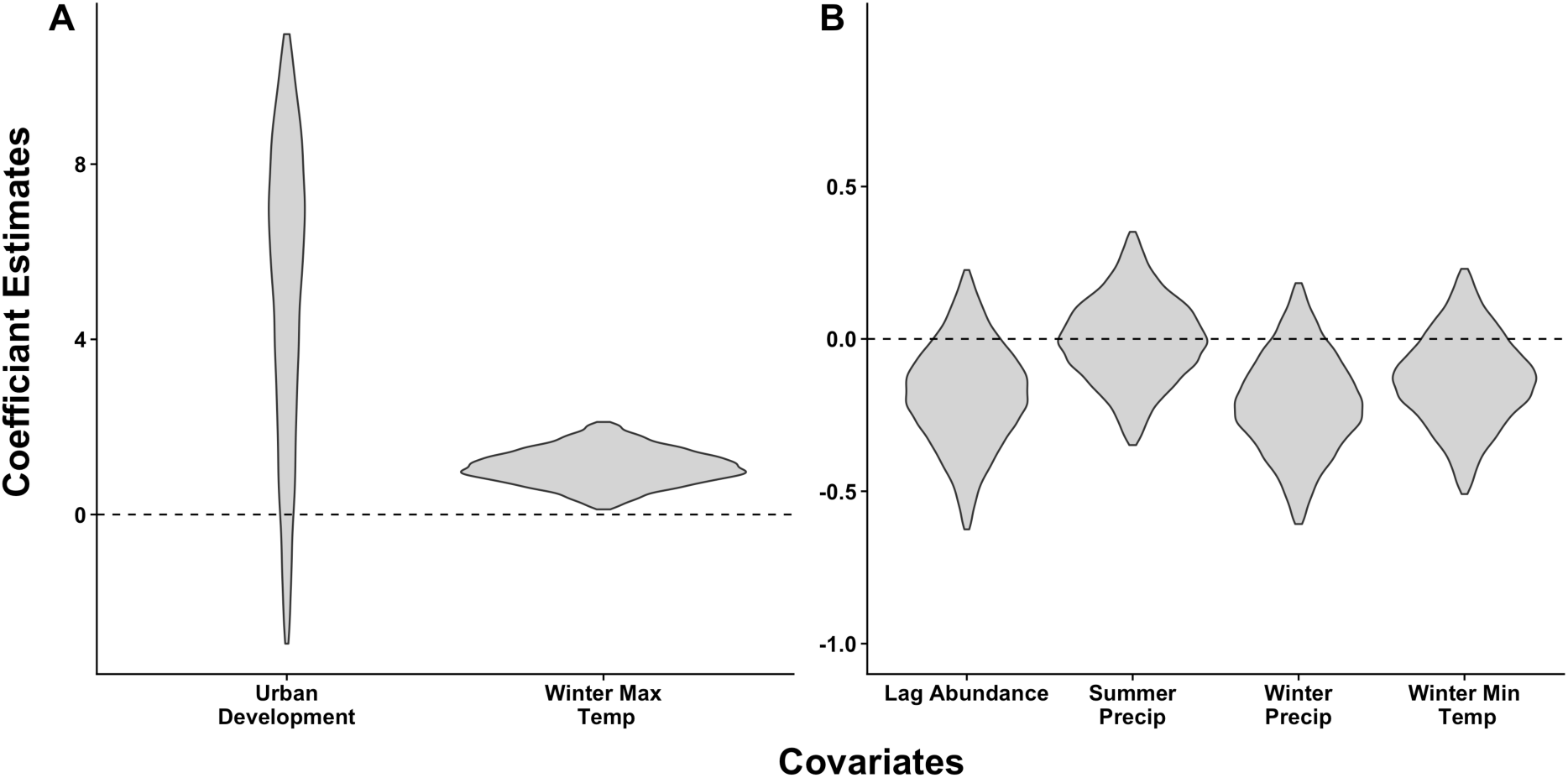
Bayesian posterior distributions for important coefficients (as determined by random forest). Y-axis shows scaled coefficient estimates. (a) Estimates of coefficients for establishment. (b) Estimates of coefficients for population growth.

### Species Distribution Models

Overall, the geographic distribution of *Passiflora* is best predicted by mean annual temperature and human population density, with the former being the most important variable (Table S2, Table 1). When examining regions (eastern and western US) separately, urban density is a more important predictor in the west, while minimum temperature is more important in the east (Table 1). All models, both combined and regional, achieved high AUC values and performed exceptionally well when compared to permuted null models (Table 1, fig. S5). Under the RCP 4.5 and 8.5 scenarios, suitable habitat along *Passiflora’s*, northern range boundary is predicted to expand. The expansion is especially strong in urban areas, with much of the urban mid-Atlantic and Northwest predicted to become more suitable (fig. 4 a,b; fig. S6 a,b).

**Table 1.** Variable importance and model fit of all species distribution models. Rows represent different regional models and columns are the different variables in the model. AUC (area under the curve) is the performance metric of model fit.

|  |  | Temperature | Population | Precipitation | AUC | OR | *P*-value |
| --- | --- | --- | --- | --- | --- | --- | --- |
| Host plant | Combined | 48.8 | 41.5 | 9.7 | 0.915 | 0.127 | **0.013** |
|  | East | 53.9 | 39.9 | 6.2 | 0.891 | 0.115 | **0.004** |
|  | West | 12.5 | 79.9 | 7.8 | 0.983 | 0.167 | **0.020** |
| Overwintering | Combined | 29.0 | 71.0 | N/A | 0.979 | 0.112 | **0.024** |
|  | East | 30.5 | 69.5 | N/A | 0.978 | 0.067 | **0.026** |
|  | West | 17.8 | 82.2 | N/A | 0.992 | 0.200 | **0.047** |
| Dispersal | Combined | 1.8 | 98.2 | N/A | 0.925 | 0.113 | **0.030** |
|  | East | 7.3 | 92.7 | N/A | 0.925 | 0.066 | **0.025** |
|  | West | 4.9 | 95.1 | N/A | 0.985 | 0.061 | 0.057 |

**Figure 4.**
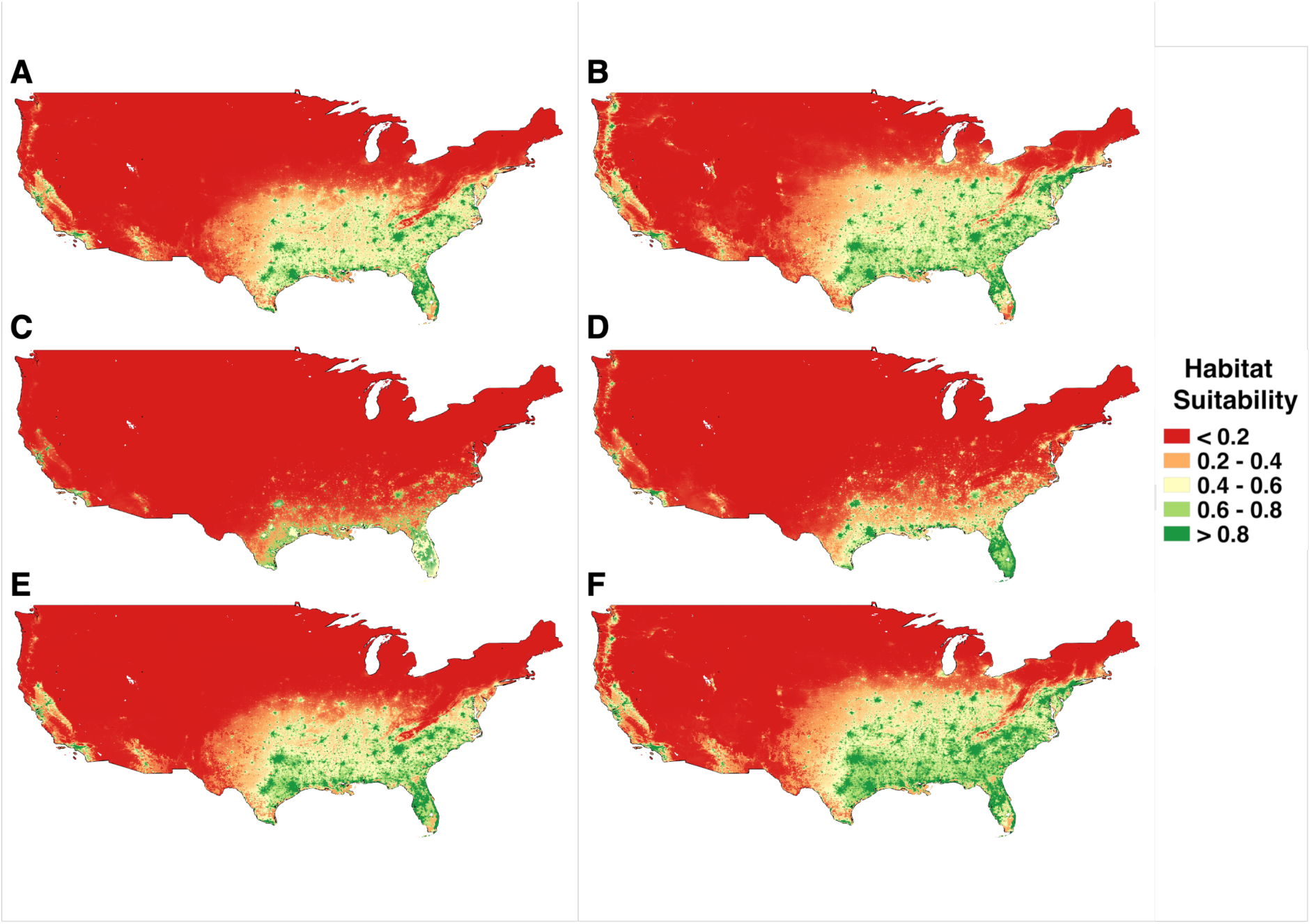
The expanding gulf fritness landscape. (a) Current distribution of suitability for *Passiflora*. (b) 2050 distribution of suitability for *Passiflora* under RCP 4.5. (c) Current distribution of suitability for overwintering *A. vanillae*. (d) 2050 distribution of suitability for overwintering *A. vanillae* under RCP 4.5. (e) Current distribution of suitability for dispersal *A. vanillae*. (f) 2050 distribution of suitability for dispersal *A. vanillae* under RCP 4.5.

The current overwintering range of *A. vanillae* is primarily in Florida and Texas, and is best explained by both *Passiflora* and winter temperature lows, as can be seen in the best-performing model with minimum temperature (Table S2). Like the host plant model, all models performed well in regards to AUC scores and in comparison to permuted null models (Table 1, fig. S5). The importance of minimum temperature in the east is greater, however in both regions host plant is more important, although it is not clear if the differences in variable importance between the east and west are meaningful given different sample sizes in the two areas (Table 1). Future climate scenarios project a slight increase in the suitability of some areas in the southeast for overwintering, but not a major expansion into new urban areas (fig. 4 c,d). The results from the models of dispersal distribution tell a different story. Models for dispersal range using different temperature variables all performed equally well, with little importance being attributed to any temperature variable in any model (Table S2). Again, models performed well using both the AUC metric and permuted null model comparison (Table 1, fig. S5). The dispersal range of the butterfly is almost entirely limited by the host plant (Table 1). This is reflected in the fact that models predict expansion in areas that closely match areas of *Passiflora* expansion (fig. 4 e,f). Thus while overwintering gains appear marginal under future warming, expansion of the range during dispersal in the summer is potentially substantial. Projections under RCP 8.5 show a slightly greater expansion, however do not dramatically vary from RCP 4.5 predictions (fig. S6 e,f).

## Discussion

Species are currently encountering novel biotic and abiotic conditions, which can positively or negatively impact population dynamics and geographic distributions (McKinney & Lockwood, 1999). Building models that parse these various stressors furthers our understanding of these impacts and allows for better prediction of future assemblages. In this study, we found that years in which the butterfly had colonized our focal sites were characterized by warmer winter maximum monthly temperatures, while winter minimum temperatures had a negative impact on population growth rates in the years after colonization. In particular, if the previous winter was cooler and drier the butterfly was found in higher abundance the next year. It is possible that the negative impact of winter climate on *A. vanillae* that we have observed is mediated through interactions with host plants or other insects. It could be the case that warmer and wetter winters negatively impact *Passiflora*, but another and perhaps more likely explanation is that wetter and warmer winters increases parasitoid pressure and/or disease leading to reduced adult emergence the following year (Harvell et al., 2002; Stireman et al., 2005). Additionally, *A. vanillae* is known to host nucleopolyhedrovirus (Rodriguez et al., 2011), which could be one mechanism that generated the observed negative density dependence (fig. S1), however this is not known to impact California populations. Finally, at our focal sites there is a slight positive trend over time in winter precipitation and winter minimum temperature (fig. S7), suggesting that if anything the butterfly is persisting and expanding in the Sacramento Valley despite of climate, not because of it.

The local impact of climate on the population dynamics of *Agraulis vanillae* in the Sacramento Valley also has implications for explaining the limiting factors for its current distribution in the west. The western United States species distribution model places almost all of the variable importance on the distribution of the host plant. One explanation for the recent colonization of the area by the butterfly is thus the increasing urbanization of the Sacramento Valley. Over the past twenty years the suburbs of Sacramento have expanded at a steady rate (Forister et al., 2010), which has likely resulted in an increase in *Passiflora* in the region. Random forest analysis ranked urban land cover over climate when predicting colonization and the Bayesian model found a much greater effect of urbanization (fig. 2a; fig. 3). In the eastern United States, the impacts of temperature, specifically minimum temperatures, are apparent in geographic distribution models. In the east, the distribution of *Passiflora* extends further north in the winter compared to *A. vanillae*, while in the west the overwintering distribution closely resembles that of *Passiflora*. Once the weather warms in the east, the butterflies can then expand to cover the distribution of the host plant. Thus, while minimum temperature plays an important role in the overwintering locations of the eastern gulf fritillary, its maximum extent appears to be host plant limited in both the eastern and western United States.

By understanding these current limits on *A. vanillae*, it is clear that any major expansion in geographic distribution will be the result of a host plant expansion. Models using the RCP 4.5 and 8.5 climate scenarios both predict geographic expansion for the host plant, and thus an expanding dispersal distribution for the butterfly. In particular, the host is predicted to have a greater presence in urban areas on both the east and west coasts, presumably through more frequent plantings into gardens that will become more suitable to the plant over time in a warming climate. If this occurs, the dispersal distribution of *A. vanillae* will also expand, as the butterfly currently tracks *Passiflora* very successfully. Dispersal ability may be an issue in the Pacific northwest, as major metropolitan areas are further apart, however given how far the butterfly currently disperses in the east each summer, it is likely this area will also be included in its distribution. Increasing temperatures may also impact the overwintering distribution of the butterfly, but given the impact of minimum temperature from the temporal analysis and the lack of major shifts from the SDM projections, this is much more uncertain. Although the full distribution of the butterfly does not appear to be directly limited by temperature, there is an indirect effect mediated by its host plant, which is limited by temperature. Projected rising temperatures will still have a major impact on the distribution of this butterfly through this indirect interaction.

Thus far, this butterfly is a notable example of a “winner” in the Anthropocene. While insect declines are occurring on a large scale (Hallmann et al., 2017; Lister & Garcia, 2018; Salcido et al., 2019; Sanchez-Bayo & Wyckhuys, 2019; Wepprich et al., 2019), altered conditions create opportunities for some insects to prevail. The intricacies of each success story are different; but an overarching theme of increasing temperature is playing a vital role in facilitating the distributional expansion of many of these insect winners. This has occurred directly by increasing the overwintering survival along a northern range margin for some species (Streifel et al., 2017), by increasing access to food resources for others (Raffa et al., 2013), or by increasing diet breadth (Pateman et al., 2012). In the case of the gulf fritillary, expansion has thus far been driven by human-mediated host plant propagation and future warming will allow this process to continue further north in the United States. While not all insect expansions will be due to temperature, ectotherms continue to be prime candidates for temperature driven distributional change, for better or for worse. Continuing to observe these phenomena and developing methods by which to understand them is critical. Here the combination of long-term time series data and large-scale citizen science spatial data allowed for a detailed examination of the underlying causes for such an expansion. As these types of data continue to become more widely accessible, the common themes behind insect distributional change in the Anthropocene will continue to become more apparent.

## Acknowledgements

We thank Ken Nussear for discussion about the distribution models and code for spatial thinning. Data were provided by the Butterfly and Moth Information Network and the many participants who contribute to its Butterflies and Moths of North America project. Data were also provided by iNaturalist and Calflora. MLF was supported by a Trevor James McMinn professorship.

## Contribution of authors

A.M.S. collected the Sacramento Valley observational data. J.H.T. and D.P.W. provided the climate data. C.A.H. conducted the statistical analyses. C.A.H. and M.L.F. wrote the manuscript with input from co-authors.

## References

Bohl, C.L., Kass, J.M. & Anderson, R.P. (2019) A new null model approach to quantify performance and significance for ecological niche models of species distributions. Journal of Biogeography, 46, 1101–1111.

Calflora: Information on California plants for education, research and conservation. [web application] (2014) Berkeley, California: The Calflora Database [a non-profit organization]. Available: https://www.calflora.org/ (Accessed: May 1, 2019).

Center for International Earth Science Information Network – CIESIN – Columbia University. (2018) Gridded Population of the World, Version 4 (GPWv4): Population Density, Revision 11. Palisades, NY: NASA Socioeconomic Data and Application Center (SEDAC). https://doi:.org/10.7927/H49C6VHW. (Accessed May 5, 2019),

Chen, I.C., Hill, J.K., Ohlemuller, R., Roy, D.B. & Thomas, C.D. (2011) Rapid Range Shifts of Species Associated with High Levels of Climate Warming. Science, 333, 1024–1026.

Flint, L. E., & Flint, A.L. (2012) Downscaling future climate scenarios to fine scales for hydrologic and ecological modeling and analysis. Ecological Processes 1:1–15.

Flint, L. E., Flint A.L., Thorne J. H. & Boynton, R. (2013) Fine-scale hydrologic modeling for regional landscape applications: the California Basin Characterization Model development and performance. Ecological Processes 2:1–21.

Forister, M.L., McCall, A.C., Sanders, N.J., Fordyce, J.A., Thorne, J.H., Obrien, J., Waetjen, D.P. & Shapiro, A.M. (2010) Compounded effects of climate change and habitat alteration shift patterns of butterfly diversity. Proceedings of the National Academy of Sciences, 107, 2088–2092.

Graves, S.D. & Shapiro, A.M. (2003) Exotics as host plants of the California butterfly fauna. Biological Conservation, 110, 413–433.

Gremillion, K.J. The Development of a mutualistic relationship between humans and maypops (*Passiflora incarntata L*.) in the Southeastern United States. Journal of Enthnobiology, 9, 135–155.

Hallmann, C.A., Sorg, M., Jongejans, E., Siepel, H., Hofland, N., Schwan, H., Stenmans, W., Muller, A., Surnser, H., Horren, T., Goulson, D. & de Kroon, H. (2017) More than 75 percent decline over 27 years in total flying insect biomass in protected areas. PLoS One, 12, e0185809.

Harvell, C.D., Mitchell, C.E., Ward, J.R., Altizer, S., Dobson, A.P., Ostfeld, R.S. & Samuel, M.D. (2002) Climate Warming and Disease Risks for Terrestrial and Marine Biota. Science, 296, 2158 – 2162.

Hijmans, R.J., Cameron, S.E., Parra, J.L., Jones, P.G. & Jarvis, A. (2005) Very high resolution interpolated climate surfaces for global land areas. International Journal of Climatology 25: 1965–1978.

Hijmans, R.J., Phillips, S., Leathwick, J. & Elith, J. (2013) dismo: Species distribution modeling. R package version 0.8-17.

iNaturalist. Available from https://www.inaturalist.org. Accessed [2019-05-01].

Kellner, K. (2019) jagsUI: A Wrapper Around ‘rjags’ to Streamline ‘JAGS’ Analyses. R package version 1.5.1.

Lister, B.C. & Garcia, A. (2018) Climate-driven declines in arthropod abundance restructure a rainforest food web. Proceedings of the National Academy of Sciences, 115, 10397–10406.

Lotts, K. & Naberhaus, T coordinators. (2017) Butterflies and Moths of North America. Data set accessed (or exported) 2019-06-13 at http://www.butterfliesandmoths.org/.

May, P.G. (1992) Flower Selection and the Dynamics of Lipid Reserve in Two Nectarivorous Butterflies. Ecology, 73, 2181–2191.

McGuire, M. (1999) Passiflora incarnata (Passifloraceae): A New Fruit Crop. Economic Botany, 53, 161–167.

McKinney M.L. & Lockwood J.L. (1999) Biotic homogenization: a few winners replacing many losers in the next mass extinction. Trends in Ecology and Evolution, 14, 450–453.

Nice, C.C., Forister, M.L., Harrison, J.G., Gompert, Z., Fordyce, J.A., Thorne, J.H., Waetjen, D.P. & Shapiro, A.M. (2019) Extreme heterogeneity of population response to climatic variation and the limits of prediction. Global Change Biology, 2019, 1–10.

Parmesan, C. (2006) Ecological and Evolutionary Responses to Recent Climate Change. Annual Review of Ecology, Evolution, and Systematics, 37, 637–669.

Parmesan, C., Ryrholm, N., Stefanescu, C., Hill, J.K., Thomas, C.D., Descimon, H., Huntley, B., Kaila, L., Kullberg, J., Tammaru, T., Tennent, W.J., Thomas, J.A. & Warren, M. (1999) Poleward shifts in geographical ranges of butterfly species associated with regional warming. Nature, 339, 579–583.

Pateman, R.M., Hill, J.K., Roy, D.B., Fox, R. & Thomas, C.D. (2012) Temperature-Dependent Alterations in Host Use Drive Rapid Range Expansion in a Butterfly. Science, 336, 1028–1030.

Powell, J.A., Russell, P., Russell, S. & Sperling, F.A.H. (2000) Northward expansion of two mint-feeding species of *Pyrausta* in California (Lepidoptera: Pyraloidea: Crambidae). Holarctic Lepidoptera, 7, 55–58.

R Development Core Team. (2013) R: A language and environmentfor statistical computing. R Foundation for Statistical Computing, Vienna, Austria. [WWW document]. URL http://www.R-project.org/ [accessed on 1 August 2019].

RColorBrewer, S., Liaw, M.A. (2018) randomForest: Breiman and Cutler’s Random Forests for Classification and Regression. R package version 4.6-14.

Raffa, K.F., Powell, E.N. & Townsend, P.A. (2013) Temperature-driven range expansion of an irruptive insect heightened by weakly coevolved plant defenses. Proceedings of the National Academy of Sciences, 110, 2193–2198.

Rodriguez, V.A., Belaich, M.N., Gomez, D.L.M., Sciocco-Cap, A. & Ghiringhelli, P.D. (2011) Identification of nucleopolyhedrovirus that infect Nymphalid butterflies *Agraulis vanillae* and *Dione juno*. Journal of Invertebrate Pathology, 106, 255–262.

Runquist, E.B., Forister, M.L. & Shapiro, A.M. (2012) Phylogeography at large spatial scales: incongruent patterns of population structure and demography of Pan-American butterflies associated with weedy habitats. Journal of Biogeography, 39, 382–396.

Salcido, D. M., Forister, M., Lopez, H. G. & Dyer, L. A. (2019) Ecosystem services at risk from declining taxonomic and interaction diversity in a tropical forest. bioRxiv, 631028.

Sanchez-Bayo, F. & Wyckhuys, K.A.G. (2019) Worldwide decline of the entomofauna: A review of its drivers. Biological Conservation, 232, 8–27.

Scott, J.A. 1986. The butterflies of North America: A natural history and field guide.

Shapiro, A.M. (2009) The Neo-Riparian butterfly fauna of western Argentina. Journal of Research on the Lepidoptera, 41, 24–30.

Shapiro, A.M. & Manolis, T.D. (2007) Field Guide to Butterflies of the San Francisco Bay and Sacramento Valley Regions.

Sibly, R.M. & Hone, J. (2002) Population growth rate and its determinants: an overview. Philosophical Transactions of the Royal Socity B, 357, 1153-1170.

ourakav, A. (2008) Notes on the biology of the gulf fritillary *Agraulis vanillae* (Lepidoptera: Nymphalidae), in North-Central Florida. Journal of the Lepidopterists’ Society, 63, 127.

Stireman III, J.O., Dyer, L.A., Janzen, D.H., Singer, M.S., Lill, J.T., Marquis, R.J., Ricklefs, R.E., Gentry, G.L., Hallwachs, W., Coley, P.D., Barone, J.A., Greeney, H.F., Connahs, H., Barbosa, P., Morais, H.C. & Diniz, I.R. (2005) Climatic unpredictability and parasitism of caterpillars: Implications of global warming. Proceedings of the National Academy of Sciences, 102, 17384–17387.

Streifel, M.A., Tobin, P.C., Kees, A.M. & Aukema, B.H. (2019). Range expansion of Lymantria dispar dispar (L.) (Lepidoptera: Erebidae) along its north-western margin in North America despite low predicted climatic suitability. Journal of Biogeography, 46, 58–69.

Thorne, J.H., Boynton, R.M., Flint, L.E. & A.L. Flint (2015) Comparing historic and future climate and hydrology for California’s watersheds using the Basin Characterization Model. Ecosphere 6(2). Online.

Walker, T.J. (1991) Butterfly migration from and to peninsular Florida. Ecological Entomology, 16, 241–252.

Warren, M.S., Hill, J.K., Thomas, J.A., Asher, J., Fox, R., Huntley, B., Roy, D.B., Telfer, M.G., Jeffcoate, S., Harding, P., Willis, S.G., Greatorex-Davies, J.N., Moss, D., Thomas, C.D. (2001) Rapid responses of British butterflies to opposing forces of climate and habitat change. Nature, 414. 65–69.

Wepprich, T., Adrion, J.R., Ries, L., Wiedmann, J. & Haddad, N.M. (2019) Butterfly abundance declines over 20 years of systematic monitoring in Ohio, USA. Plos ONE, 14, e0216270.

