## Supplemental Tables and Figures for "A winner in the Anthropocene: changing host plant distribution explains geographic range expansion in the gulf fritillary butterfly"

Contents:

Supporting Information Table S1: Details on spatial thinning

Supporting Information Table S2: Spatial model scores for various temperature variables

Supporting Information Figure S1: Relationship between population growth and covariates

Supporting Information Figure S2: Correlation matrix of all model covariates

Supporting Information Figure S3: Individual site level Bayesian posterior distributions from the colonization model

Supporting Information Figure S4: Individual site level Bayesian posterior distributions from the population growth model

Supporting Information Figure S5: Real SDM comparison to permuted null models.

Supporting Information Figure S6: *Passiflora, A. vanillae* overwintering, and *A. vanillae* migratory SDMs under RCP 8.5

Supporting Information Figure S7: Change in winter minimum temperature and winter precipitation over time.

Table S1. Number of observations before and after thinning for each species distribution model.

| | | Total Points | After Thinning |
|---|---|---|---|
| Host plant | Combined | 1606 | 894 |
| | East | 1532 | 823 |
| | West | 83 | 71 |
| Overwintering | Combined | 571 | 327 |
| | East | 437 | 247 |
| | West | 134 | 68 |
| Migratory | Combined | 4972 | 1743 |
| | East | 4169 | 1535 |
| | West | 803 | 208 |

Table S2. AUC scores for Maxent models of with various temperature inputs.

| | | AUC | Importance |
|---|---|---|---|
| Host plant | Average Temperature | 0.925 | 47.7 |
| | Max Temperature | 0.916 | 42.7 |
| | Min Temperature | 0.909 | 50.4 |
| Overwintering | Average Temperature | 0.951 | 21.5 |
| | Max Temperature | 0.970 | 8.8 |
| | Min Temperature | 0.982 | 36.2 |
| Migratory | Average Temperature | 0.925 | 2.4 |
| | Max Temperature | 0.924 | 2.9 |
| | Min Temperature | 0.926 | 5.7 |

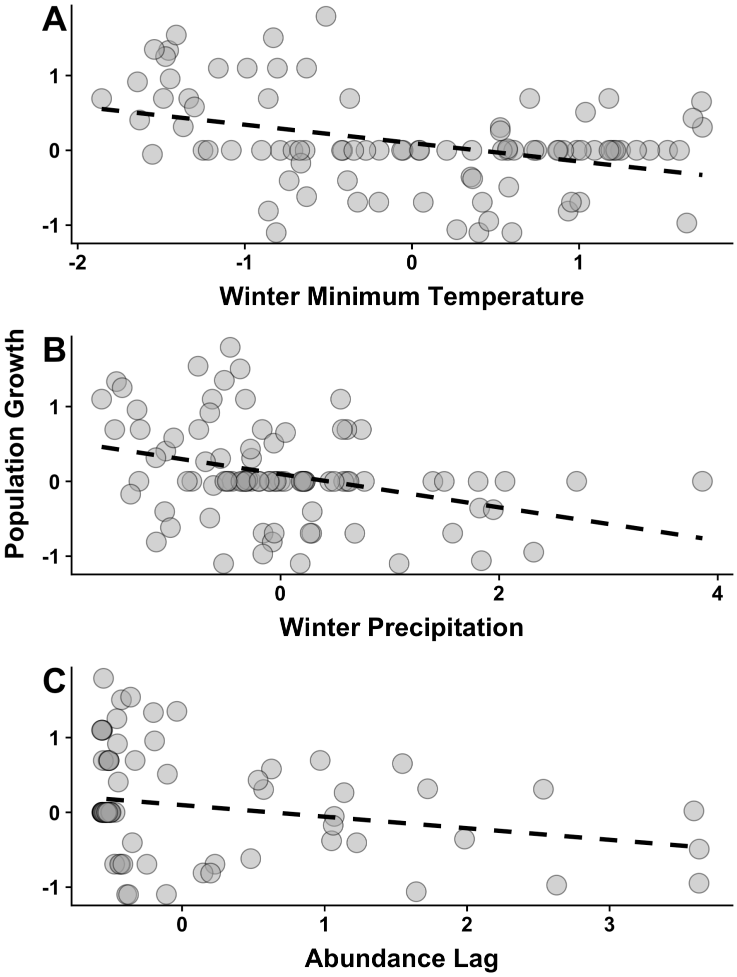


Figure S1. (a) Relationship between population growth and winter minimum temperature in the previous year. (b) Relationship between population growth and winter precipitation in the current year. (c) Relationship between population growth and butterfly abundance in the previous year. Points are from across all sites.

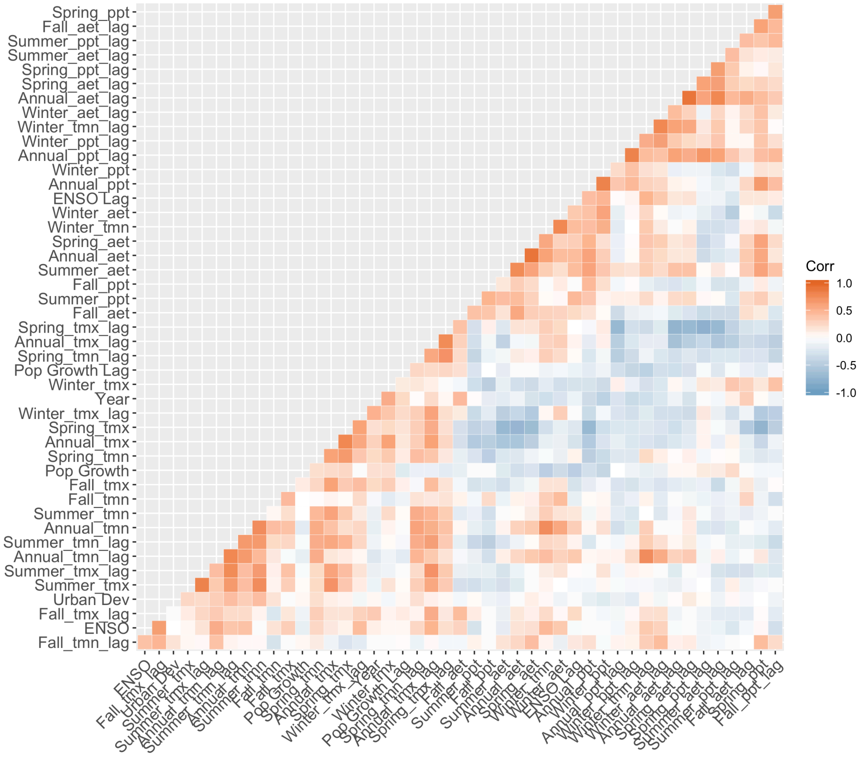


Figure S2. Correlation matrix of all climate variables, lagged climate variables, year, urban development, and population growth.

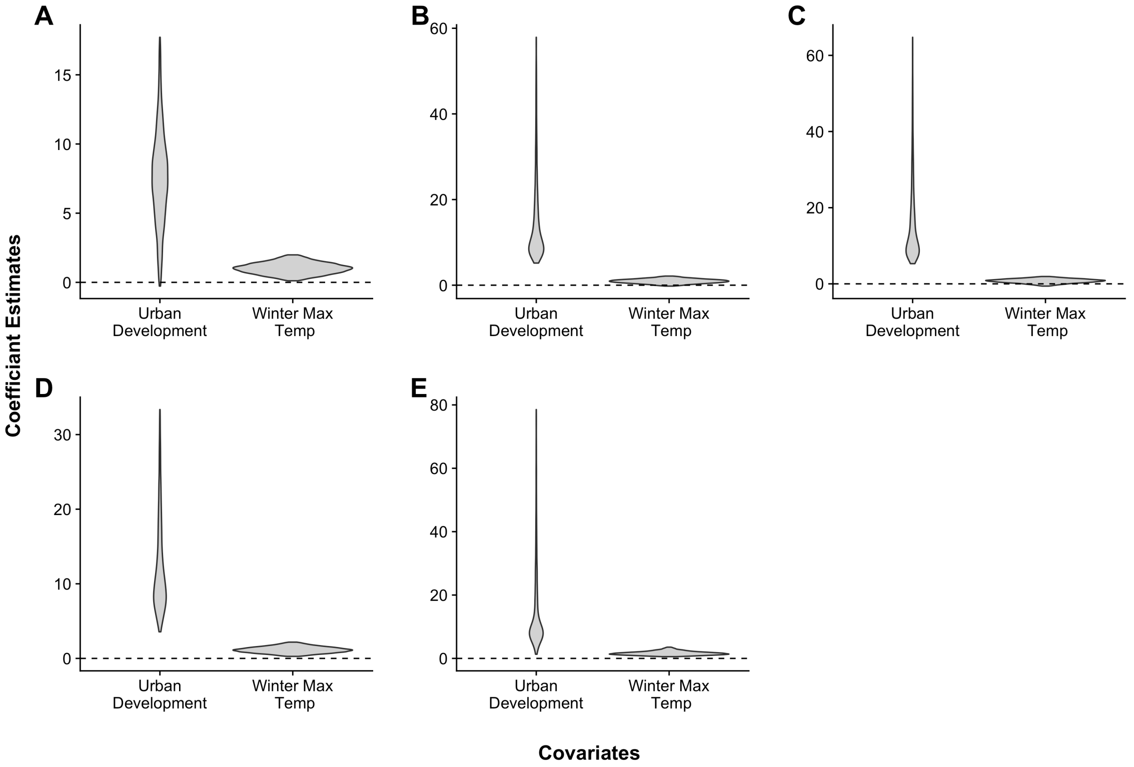


Figure S3. Individual site level Bayesian posterior distributions from the colonization model. Y-axis shows scaled coefficient estimates. (a) Gates Canyon (b) North Sacramento (c) Rancho Cordova (d) Suisun Marsh (e) West Sacramento.

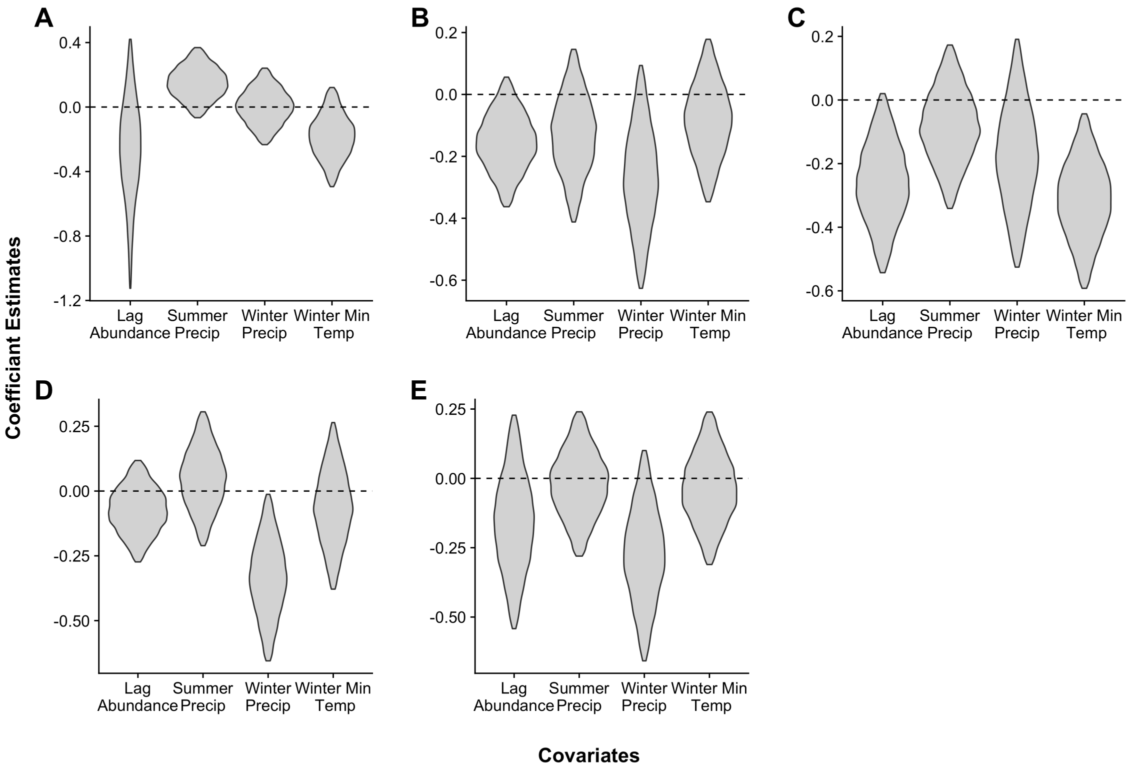


Figure S4. Individual site level Bayesian posterior distributions from the population growth model. Y-axis shows scaled coefficient estimates. (a) Gates Canyon (b) North Sacramento (c) Rancho Cordova (d) Suisun Marsh (e) West Sacramento

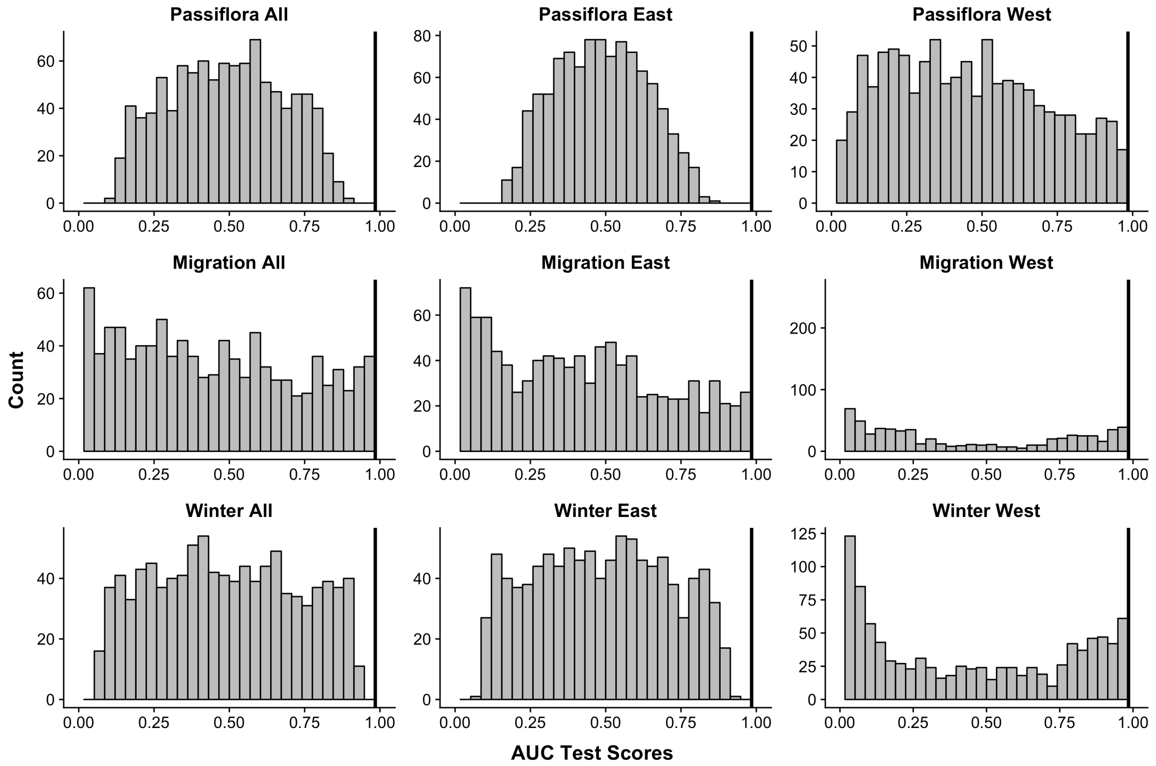


Figure S5. Comparison of SDMs to permuted null models. The solid black line shows the AUC score of the true model compared to a histogram of null model AUC scores. AUC scores are the scores of the testing data.

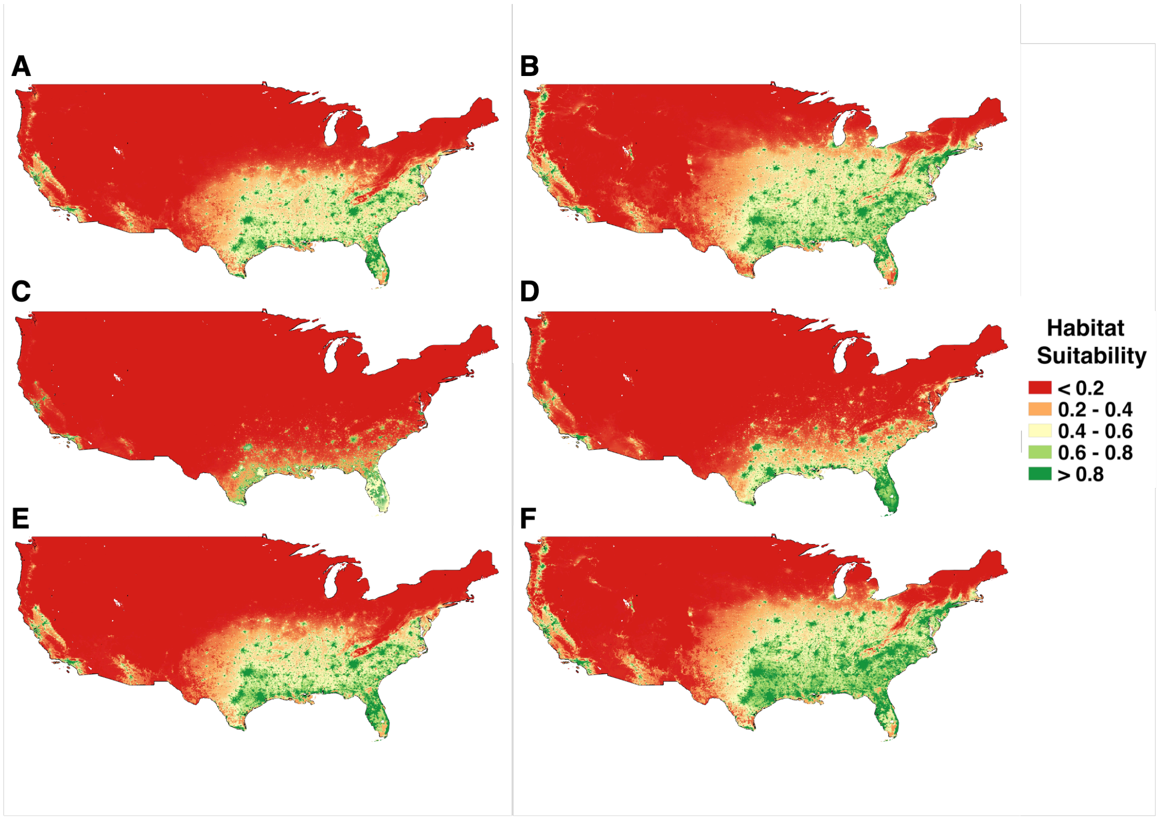


Figure S6. (a) Current distribution of suitability for *Passiflora*. (b) 2050 distribution of suitability for *Passiflora* under RCP 8.5. (c) Current distribution of suitability for overwintering *A. vanillae*. (d) 2050 distribution of suitability for overwintering *A. vanillae* under RCP 8.5. (e) Current distribution of suitability for migratory *A. vanillae*. (f) 2050 distribution of suitability for migratory *A. vanillae* under RCP 8.5.

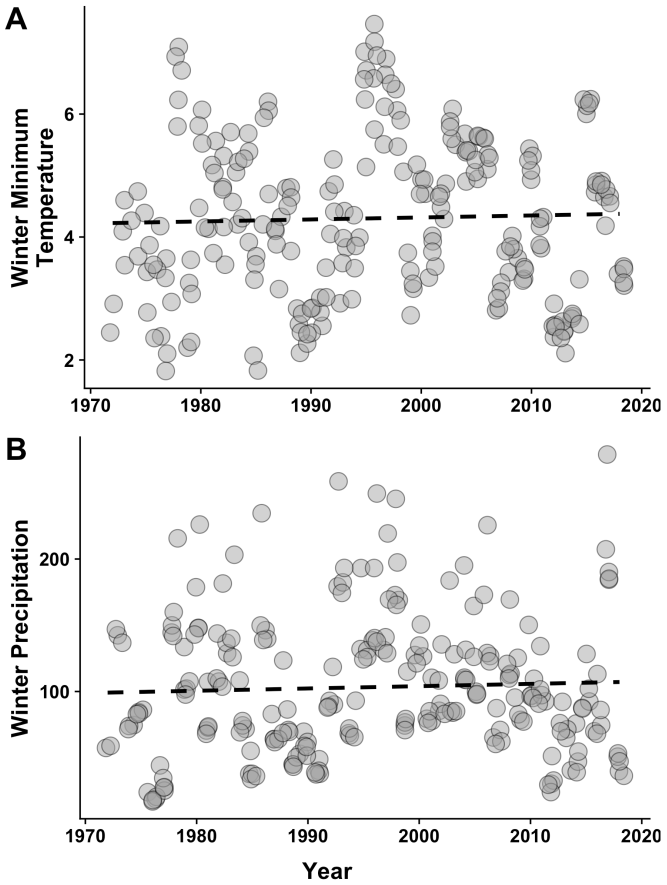


Figure S7. (a) Relationship between winter minimum temperatures and time. (b) Relationship between winter precipitation and time. Points are from across all sites.
